# Kinetic analysis of CSF–brain tracer exchange in the pig brain under different anesthetic regimes

**DOI:** 10.64898/2026.08.24.746655

**Authors:** Miriam L. Navarro, Anders S. Olsen, Sara M. Ulv Larsen, Clara Madsen, Robin de Nijs, Katarina Bubulovic, Julius Søndergaard, Cyril Pernet, Louise Møller Jørgensen, Claus Svarer, Gitte Moos Knudsen

## Abstract

**Introduction:** Anesthesia is known to modulate glymphatic clearance and cerebrospinal fluid (CSF) transport in rodents, but how these effects translate to a larger, gyrencephalic brain is unknown. With its anatomical similarity to the human brain, the pig offers a valuable translational model for examining anesthesia-dependent CSF-to-brain transport.

**Methods:** We used dynamic *in vivo* SPECT/CT imaging for six hours following cisterna magna injection of [99mTc]-DTPA to quantify CSF-to-brain tracer transport in pigs under two anesthesia regimens: ketamine/dexmedetomidine (K/D, n=5) which previously has been shown in rodents to enhance glymphatic influx relative to GABAergic anesthesia, and propofol (PRO, n=5). Brain and CSF spaces were delineated using a data-driven non-negative matrix factorization approach, and tracer kinetics were quantified using a one-tissue compartment model.

**Results:** Brain influx could be stably estimated from 2 hours post-injection. Hierarchical sub-division of the brain parenchyma identified two kinetically distinct components with different anatomical distributions: a surface component, located ventrally and within the interhemispheric fissure, showed faster kinetics than the anatomically deeper and lateral-dorsal component. Consistent with rodent findings, K/D-anesthetized pigs showed 62% (p=0.002) greater brain tracer accumulation than PRO-anesthetized pigs. However, while the brain influx rates did not differ substantially (p=0.047), a 52% higher cumulative CSF tracer concentration (p=0.047) could account for most of the difference by providing greater tracer availability for brain entry.

**Conclusions:** In the larger gyrencephalic pig brain, we found higher brain tracer accumulation under K/D anesthesia compared to PRO anesthesia. A significant portion of this difference is readily explained by higher CSF retention, likely driven by a slower CSF turnover. This underscores the necessity of dynamic CSF tracer concentration measurements when assessing CSF-brain influx, a factor we suggest that future glymphatic studies should take into account.

## 1. Introduction

The identification of a coordinated flow of cerebrospinal fluid (CSF) through perivascular and paravascular spaces into the brain parenchyma, often referred to as the glymphatic system^1,2^, has become a growing focus in neuroscience. Rodent studies have been instrumental in this research where it has been demonstrated that glymphatic activity fluctuates with brain states such as sleep^3–6^ and anesthesia^7^. Sleep-like anesthetics such as ketamine combined with α_2_-adrenergic drugs like dexmedetomidine (DEX) or xylazine have been repeatedly reported to enhance tracer influx from CSF into the brain parenchyma compared to GABAergic anesthetics such as isoflurane^7–13^ or propofol (PRO)^14^. The mechanism underlying the DEX-associated increased influx has been ascribed to decreased central noradrenaline-driven vasomotion or tone in the locus coeruleus^15^, aquaporin-4 polarization^16^, reduced intracerebral blood volume^17^, and delta-wave activity^7^ as drivers of CSF flow within the brain parenchyma.

Perioperative neurological complications such as delirium or postoperative cognitive dysfunction are often seen in particularly elderly patients and after prolonged anesthesia exposure^18–20^. Given that there is some evidence that either peri- or postoperative administration of DEX can ameliorate patients’ condition^21–24^, it has been speculated that these complications may originate from reduced brain waste clearance. So far, most studies in humans aiming at exploring CSF dynamics have been based on MRI-based approaches, such as measuring CSF flow at the aqueduct or fourth ventricle^25–27^ or administering intrathecal contrast agents to assess their distribution and clearance^28–30^. These clinical studies have measured CSF dynamics and tracer CSF clearance from the parenchyma, but not at a temporal resolution that allows for full kinetic analysis that quantifies tracer brain influx and efflux.

Importantly, the rodent lissencephalic brain may differ from gyrencephalic brains in its brain fluid organization which can make it difficult to translate findings from rodents to humans. Large-animal models offer a route to bridging this gap. The porcine brain shares key features with the human brain, including gyrencephaly, a high white-to-grey matter ratio, comparable size, and diurnal sleep-wake cycles^31–35^, making it an increasingly used neuroscience model^35–41^. We have previously presented the porcine model for assessment of the glymphatic system, including detailed methodology for intrathecal tracer injection and dynamic acquisition of Single Photon Emission Computed Tomography (SPECT)^14^.

Here, we use dynamic *in vivo* combined SPECT and x-ray Computed Tomography (CT) imaging over six hours following cisterna magna injection of [^99m^Tc]-DTPA to quantify CSF-to-brain tracer transport in pigs under PRO or K/D anesthesia, adopting an automatic data-driven delineation approach using non-negative matrix factorization (NMF) followed by one-tissue compartment modelling (1TCM). Moreover, detailed physiological monitoring (vitals, ICP, EEG), and whole-body tracer distribution and excretion enable us to qualify the effects of the anesthetic regimens.

## 2. Methods

### 2.1 Animals and anesthesia

All experiments were approved by the Danish Veterinary and Food Administration’s Council for Animal Experimentation (license No. 2022-15-0201-01156), and followed the ARRIVE guidelines^42^ and were conducted in accordance with the European Communities Council Resolves of 22 September 2010 (2010/63/EU).

We used ten female pigs weighing around 20 kg (∼10 weeks-old) (crossbred Landrace, Yorkshire and Duroc); pig care and preparation conditions are described in our previous publication^14^. Here, we extend the analysis to include a scanning time from three hours to six hours and instead of manually define regions of interest (ROIs) and compare across anesthesias only using areas under the curve (AUCs), we describe an automatic brain segmentation method and compare anesthesias also with kinetic modeling. In this study we also analyzed intracranial pressure (ICP) and cortical activity by encephalography (EEG).

All experiments were performed during the light phase of the animals, equivalent to the awake stage for pigs. On the experimental day, pigs were premedicated by intramuscular injection of 0.14 mL/kg Zoletil® 50 Vet (Virbac, Kolding, Denmark) mixture, weighed, and intubated. Two anesthesia regimens were applied: (1) PRO group: maintenance anesthesia via intravenous infusion of propofol (12.5 mg/kg/h; Fresenius Kabi AS, Halden, Norway) and fentanyl (5 µg/kg/h, Fentanyl 50 µg/mL, Hameln, Germany) for 6 hours for all animals (PRO1-5); or (2) K/D group: an initial intramuscular dexmedetomidine (DEX) (Dexmedetomidinehydrochlorid 0,5 mg/ml, Orion Pharma, Finland) dose (18 µg/kg for KD1–KD3; 150-200 µg/kg for KD4–KD5 pigs), followed by continuous DEX infusion (8–16 µg/kg/h) and ketamine infusion (14 mg/kg/h, Ketaminol 100 mg/ml, Vetviva, Austria) throughout scanning, plus a second intramuscular DEX dose administered at ∼3 hours post-injection; 150–200 µg/kg for KD1–KD3, and 80 µg/kg and 45 µg/kg for KD4 and KD5, respectively. The second dose was decided in order to maintain the pigs under anesthesias for six hours, while the dosage difference across K/D group was a result from a change in the experimental protocol half-way through the study. Despite the dosage differences, all K/D pigs had a similar cumulative DEX dose over the scanning period. In addition, compared to KD1-3 pigs, KD4 and KD5 did not differ in delta wave proportion, cerebral perfusion pressure (CPP), or heart rate (Figure 1C and supplementary Table S1).

**Figure 1.**
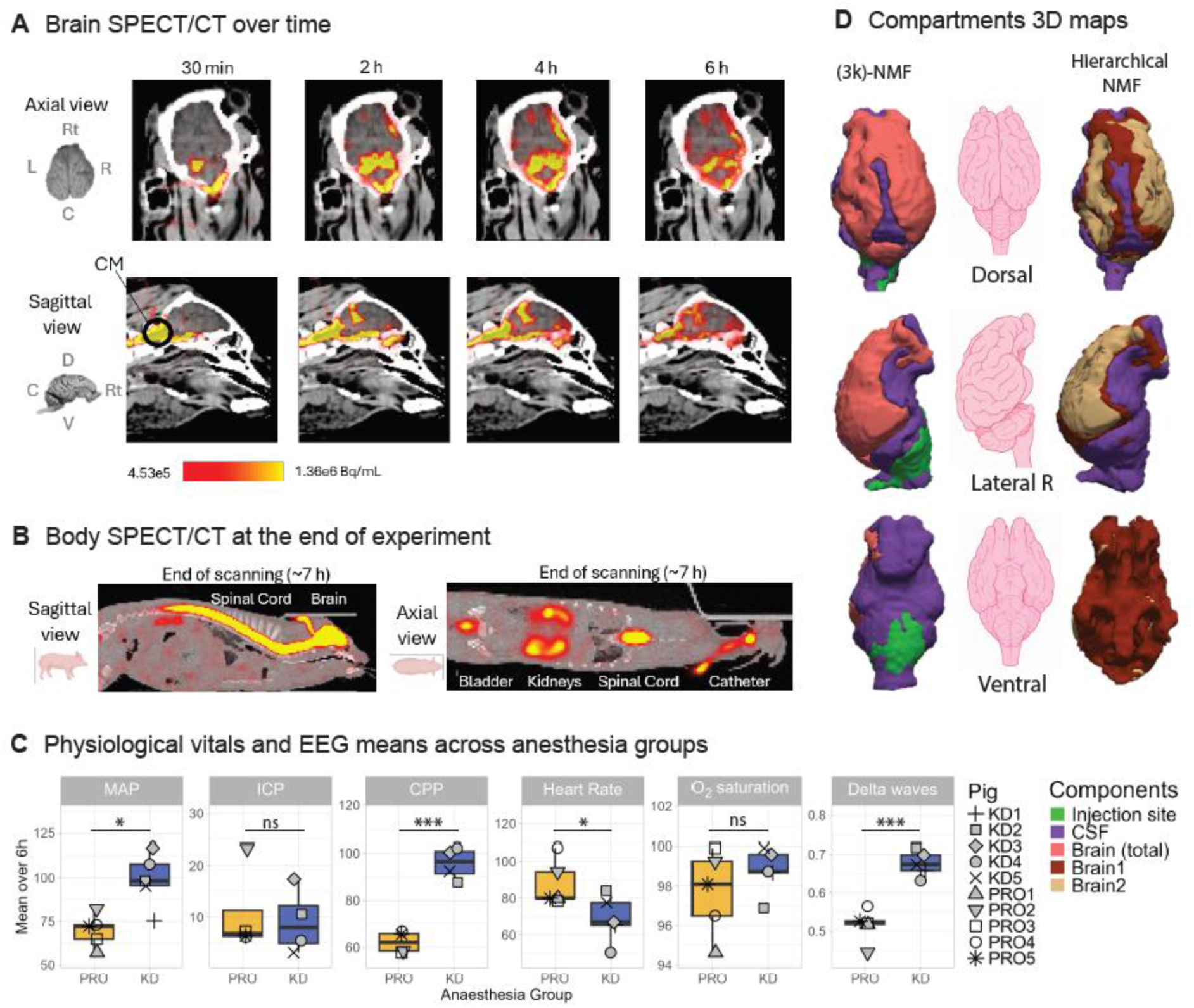
Dynamic SPECT/CT images, vitals and EEG characteristics, and component maps. (A) Representative axial (top) and sagittal (bottom) SPECT/CT images showing [^99m^Tc]-DTPA tracer distribution in the brain over the 6-hour dynamic scan (30 min, 2 h, 4 h, and 6 h post-injection). The cisterna magna (CM) injection site is indicated on the sagittal view. Color scale represents radioactivity concentration. Anatomical orientation is indicated (Rt = right, L = left, C = caudal, D = dorsal, V = ventral). (B) Whole-body SPECT/CT images from the end of the experiment (∼7 hours post-injection), showing sagittal (left) and axial (right) views of tracer distribution across the spinal cord, brain, bladder, kidneys, and the CM catheter. (C) Physiological vitals and EEG delta power, averaged across the full 6-hour scanning period and compared between anesthesia groups (PRO, propofol; KD, ketamine/dexmedetomidine). Boxplots show mean arterial pressure (MAP), intracranial pressure (ICP), cerebral perfusion pressure (CPP), heart rate, oxygen saturation (O₂ saturation), and EEG delta wave power. Individual points represent individual pigs (shapes as indicated in the legend: KD1–KD5, PRO1–PRO5). Asterisks indicate statistical significance from unpaired t-tests (* p < 0.05, *** p < 0.001, ns = not significant). (D) Three-dimensional reconstructions of brain and CSF components derived from non-negative matrix factorization (NMF), shown in dorsal, lateral (right), and ventral views. Left column: (3k)-NMF delineation of the whole intracranial space (injection site green, CSF in purple, brain parenchyma in pink). Middle column: reference pig brain for anatomical orientation. Right column: hierarchical NMF sub-division of the brain parenchyma (Brain1 in dark-red and Brain2 in beige) and CSF (purple).

### 2.2 Experimental setting and protocol

Pigs were monitored throughout the SPECT-scan by EEG, visual inspection, blink reflexes, and manual and digital recording of mean arterial pressure (MAP), heart rate (HR), end-tidal CO_2_ (EtCO_2_), oxygen saturation (SpO_2_), and temperature were collected during the scanning period as described in our previous publication^14^. We also measured ICP from the CM catheter connected to a Flushless TruWave^TM^ transducer (Edwards Lifesciences Corporation, California, USA) after the tracer infusion and throughout the experiment. EEG was also measured throughout the scan, described in the following section. Blood partial CO_2_ (pCO_2_), blood pH, glucose and lactate levels were measured in arterial blood samples collected at discrete time points, stored on ice, and analyzed in an ABL90 FLEX (Radiometer, Denmark) every 3-4h^43,44^. Glucose and saline were infused during the scanning as needed, based on the animal’s needs. Respiration frequency and volume were controlled by a mechanical ventilator with ∼34% O_2_. Digital recording of monitor’s vitals was done using VitalRecorder^45^ software. CPP is calculated subtracting the ICP from the MAP. Pulse pressure is the difference between systolic and diastolic. Physiological vitals are presented in Table S1 as mean and standard deviation (SD) over the six hours of scanning for each pig.

The CM cannulation was done with the pig in left lateral position and to mimic a comfortable natural sleeping position of the pig, this position was maintained during the scanning. 1mL [Tc-99m]-DTPA (12.5 mg/mL, TechneScan DTPA, Curium Pharma; MW 489 Da prepared at the Copenhagen University Hospital -Rigshospitalet) at a rate of 0.05 mL/min and flushed with 0.1 mL of artificial CSF. SPECT scanning was started simultaneously with the start of tracer infusion. After the six hours dynamic brain SPECT scan, a whole-body scan (SPECT and CT) was done in all but two pigs. The pig PRO1 whole body scan did not include the head and upper part of the spinal cord and due to missing personnel, pig KD5 did not get a whole-body scan; these pigs were not included in the final body radioactivity measurements. Immediately after the last scans had been acquired, pigs were euthanized with an overdose of 15-20 mL sodium pentobarbital (ScanVet Animal Health A/S, Fredensborg, Denmark).

Arterial blood samples were manually drawn at 0, 5, 10, 20 minutes and thereafter every 20 minutes post scan start. Urine samples were collected every 30 minutes from the urinary bladder catheter (draining into a collection bag); in cases where no urine had been produced, the sampling was skipped. When the urine was sampled, the total urine volume in the collection bag was noted and the urine bag was emptied. Radioactivity in whole blood and urine samples was measured with an automated gamma-counter (Hidex AMG, Hidex Oy, Turku, Finland) and decay corrected to start of the scanning.

### 2.3 Electroencephalogram

EEG was measured with 9 sub-dermal needle electrodes inserted in the top skin of the skull. The needles were placed using the midpoint between the ears as reference point: two frontal needles (8 cm from the reference point) at the left and right sides, each 1 cm from the midline; three central needles (6 cm from reference point), right and left were 1 cm from the midline needle; two parietal needles (4 cm from reference point) also right and left 1 cm apart from midline, and a reference needle in the ear skin and a ground needle in the neck skin. The needle electrodes were connected to a Cadwell Easy II amplifier (Cadwell, Washington, USA) and recorded with EEG Software (Cadwell Easy PSG Software, version 2.0.1, Washington, USA).

We computed the multitaper spectrogram^46^ of the C3-ref channel and the band power in the delta (0.5-4Hz), theta (4-8Hz), alpha (8-12Hz), beta (12-30Hz), and gamma (30-100Hz) ranges relative to the total power. Bandpowers are shown in Table S2.

### 2.4 SPECT/CT acquisition

Details on SPECT acquisition and preprocessing are described in Beschorner, Navarro et al^14^. Briefly, scans were acquired on a Mediso AnyScan Trio SPECT/CT (Mediso, Budapest, Hungary) equipped with a 16-slice diagnostic CT and three gamma camera heads with 9.5 mm thick sodium iodide crystals mounted with multi-pinhole (MPH) brain collimators. The frame duration including camera head movements was 4-6 minutes. The individual SPECT-frames were reconstructed into 1.8x1.8x1.8 mm voxel size, calibrated for ^99m^Tc and saved in Bq/mL. Figure 1A shows representative brain SPECT/CT images at different times and Figure 1B displays a representative whole-body SPECT/CT image.

### 2.5 Brain and CSF delineation using non-negative matrix factorization

To delineate brain tissue, we opted for a data-driven method using 3-component non-negative matrix factorization ((3k)-NMF)^47^ of the SPECT-data. This approach was selected to reduce the influence of partial volume mixing and radioactivity spillover between brain and CSF volumes, compared to anatomical segmentation. NMF decomposes a non-negative spatiotemporal dataset into spatial and temporal components. To prevent voxels outside the brain from influencing the decomposition, voxels with a summed radioactivity concentration below 5MBq/mL over the entire acquisition period were excluded prior to the NMF. Due to significant peak radioactivity differences between components, we normalized the time-activity curves (TACs) for each remaining voxel by the summed radioactivity counts over the entire acquisition period. This normalization was performed to make the decomposition primarily sensitive to the shape rather than the amplitude of TACs. Following NMF, the resulting spatial components were used to derive non-normalized TACs from the original voxel-wise SPECT data. In this framework, each voxel was assigned a continuous weight for each component, reflecting the degree to which the voxel belonged to the corresponding tissue component. Component-specific TACs were thus calculated as the weighted mean of the voxel TACs within each spatial component. To further reduce potential partial volume effects, voxels with weights below the 30th percentile of each spatial map were excluded prior to calculating the weighted mean TAC. The 30^th^ percentile threshold was selected to minimize spillover of radioactivity from CM into the brain parenchyma. Finally, the TACs were normalized to the injected dose and divided by the body weight of each pig resulting in a unitless “normalized radioactivity concentration”.

While a two-component decomposition mainly separated the intracranial cavity into regions spatially corresponding to the injection site (CM) and non-CM CSF spaces, a three-component decomposition also resulted in a brain-component^48^. Increasing the number of components mainly led to a finer subdivision of the CM region, likely due to the fast CSF dynamics influenced by the bolus radiotracer injection. Therefore, to further describe brain influx dynamics, we chose a hierarchical approach to subdivide the brain parenchyma into regions with different radioactivity patterns, by estimating a two-component NMF only on brain voxels. Thereby, we divided the brain parenchyma into two subcomponents (“Brain1” and “Brain2”).

We estimated the volume of each SPECT-derived component by hard-assigning each voxel to the component with the highest weight, then summing the number of assigned voxels for each component and multiplied with the voxel volume in cm^3^. For visualization purposes, we binarized the weighted NMF maps, to show three-dimensional maps.

### 2.6 Brain and body radioactivity measurements

Blood radioactivity obtained from the gamma-counter (measured in Bq/mL) was decay corrected to start of scan time and interpolated to scan times for each pig and then subtracted from the brain parenchyma TAC assuming 5% of blood in the brain. NMF component TACs were normalized to the injected dose and to the pig body weight resulting in a normalized unitless estimate of radioactivity.

For blood radioactivity, we calculated the percentage from the injected dose (%ID), multiplying the radioactivity concentration in Bq/mL by multiplying the average total blood volume in a 20 kg pig (65 mL/kg) and dividing by the injected dose. Correspondingly, urine radioactivity concentration (Bq/mL) was multiplied by the urine volume excreted (in mL) and summed over time to obtain cumulative urine radioactivity, and then the final decay-corrected cumulative urine radioactivity was divided by the injected dose.

For the whole-body tracer distribution, we manually drew ROIs in the final whole-body scans using ITK-SNAP (version 4.0.1)^49^, including: intracranial space, entire spinal cord, bladder, and kidney. The spinal cord was further subdivided in the cervical (C1-C7), thoracic (T1-T14), lumbar (L1-L6) and sacral (rest of vertebra) sections. TACs for each ROI were defined as the average of the voxels inside the ROI. Body TACs were multiplied with the ROI volume to obtain total radioactivity (MBq) which was subsequently normalized to the injected dose (%ID). Final urine radioactivity was calculated using the last cumulative urine value of the scanning period plus the bladder radioactivity.

AUC was calculated with the trapezoidal rule implemented in the *trapz* function from the *pracma* package in R (version 4.6.0)^50^, for the following TACs: NMF TACs (injection site, CSF, brain, and the hierarchical regions Brain1 and Brain2) and blood TACs, for each animal. Peak radioactivity for the injection site and CSF TACs was selected as the highest radioactivity value in the injection and CSF TACs for each pig.

### 2.7 Kinetic Modelling

We used a one-tissue compartment kinetic model (1TCM) to assess the brain tracer motion from CSF to brain (brain influx = K_1_) and back (brain efflux = k_2_), with the CSF TAC as an input function. To assess the stability of the modeling, we quantified end-truncated data by shortening the TACs by one hour (with “free”, not fixed parameters). Based on the average k_2_ (0.13 h^-1^), determined from the 6 h acquisitions for both anesthesia groups, we repeated the modeling by fixing k_2_ to either 0 h^-1^ or to the average k_2_. The kinetic modelling was done with PMOD (RRID:SCR_016547, http://www.pmod.com).

### 2.8 Generative AI

Generative AI tools (ChatGPT-4o^51^ and Claude Sonnet 3.5^52^) were employed for two purposes: (1) assisting in the development of R code for handling data, statistical analysis, and visualization of data; and (2) improving the linguistic clarity and flow of the manuscript. All AI outputs were carefully reviewed, verified, and edited by the authors. The authors assume full responsibility for the accuracy of the data, validity of the hypotheses, and integrity of all content presented in this article.

### 2.9 Statistical analysis

Unpaired Welch’s t-tests were used to compare mean physiological vitals, EEG band power, AUCs (from NMF-derived and blood TACs), kinetic modelling parameters (K_1_, k_2_, and K_1_/k_2_ = V_T_), and %ID values (body-ROIs and urine) between the two anesthesia groups. Given the six hours duration of the scanning protocol and that DEX maintenance dose was given around three hours post-injection, we additionally tested with a paired t-test whether physiological and EEG values differed between the first half of scanning (0–3 h post-injection) and the second half (3–6 h post-injection). For assessment of kinetic parameter stability over time, we used paired t-tests to compare influx rate constants (K_1_) across end-truncated datasets, with each truncated K_1_ estimate compared against the 6-hour reference value, pooling both anesthesia groups (PRO and K/D). All statistical tests were two-tailed. Statistical significance was set at 0.05. Statistical analyses were performed in R (version 4.3.1; R Core Team, 2024).

## 3. Results

### 3.1 EEG and vitals

Figure 1 summarizes key physiological vitals and EEG band power across the 6-hour scanning period for the two groups. Individual pig data are listed in the Supplementary (Table S1 and Table S2). The main physiological differences are illustrated in Figure 1C: the K/D group showed higher MAP, CPP, and delta wave power than the PRO group, alongside lower heart rate, and relative alpha wave power, with no difference in ICP or O_2_ saturation. In general, when comparing the first three hours of scanning with the last three hours, only minor differences were observed (Table S1 and S2): The PRO group showed a minor reduction of 4% in MAP and the K/D group showed an increase of 9% in systolic pressure and a 11% decrease in heart rate. Importantly, EEG delta wave power remained unchanged throughout the entire experimental period for both groups.

### 3.2 NMF component maps and TACs

In order to delineate the injection site (CM and surroundings) from CSF and the brain parenchyma, we first tested a three-components (3k)-NMF decomposition^48^ where SPECT data were decomposed based on the shape of TACs. Next, we manually classified the components based on the anatomical location into: injection site, CSF, and brain parenchyma. Fewer components failed to separate the brain parenchyma from the CSF; adding more components primarily subdivided the injection site. A representative example of a NMF 3D map is shown in Figure 1D and their TACs in Figure 2A; segmentation maps are shown for all individual pigs in Supplementary Material Figure S1.

**Figure 2.**
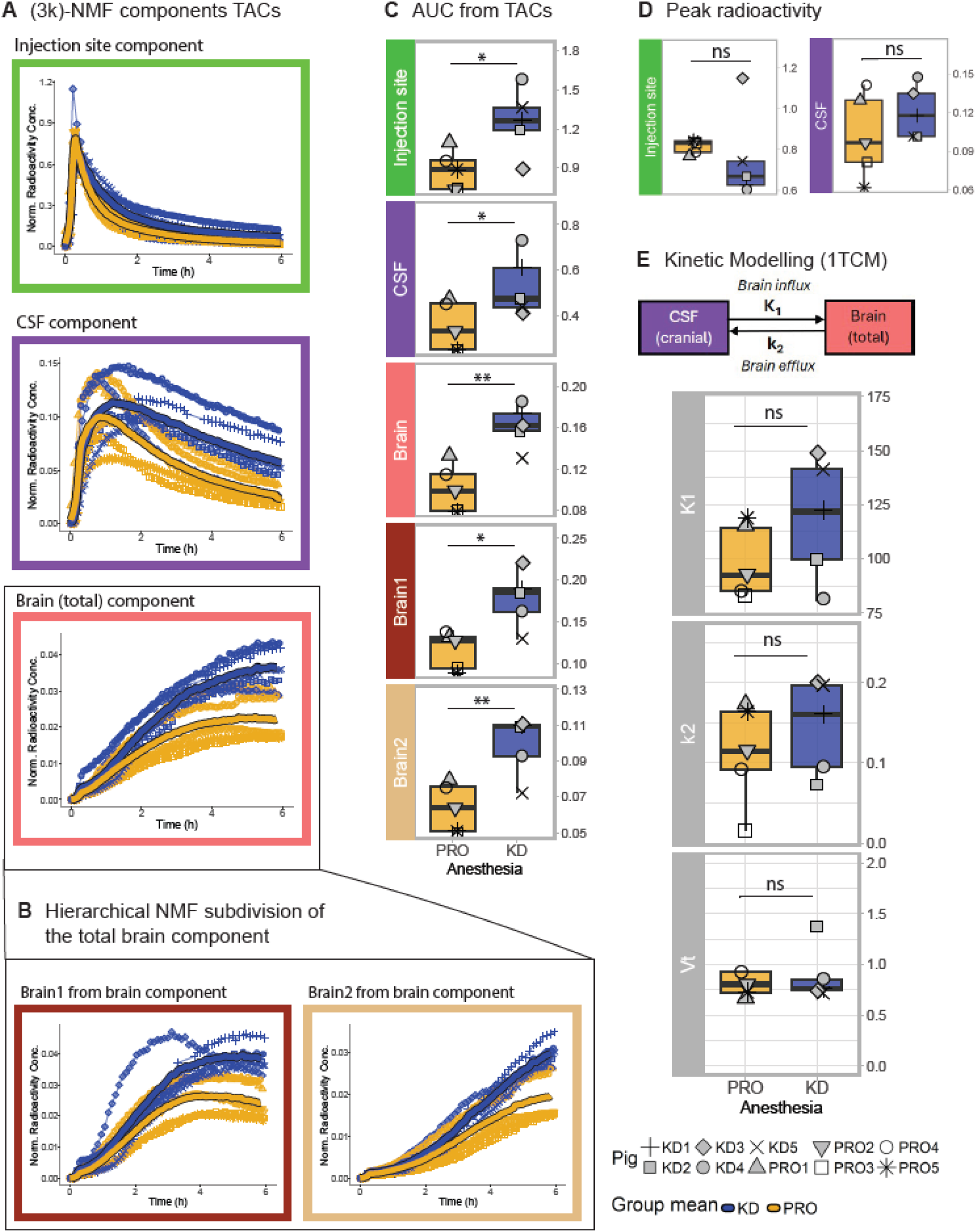
Regional time-activity curves, area under curve (AUC), peak radioactivity, and kinetic modelling parameters (K_1_, k_2_, V_T_) for CSF-to-brain tracer transport under propofol and ketamine/dexmedetomidine anesthesia. (A) Time-activity curves (TACs) for the injection site, CSF, and brain parenchyma derived from (3k)-NMF decomposition of the SPECT data, plotted per pig (thin lines with points) with group means overlaid (thick lines; blue = KD, orange = PRO). Y-axis shows normalized radioactivity concentration; x-axis shows time post-injection (hours). (B) Hierarchical NMF subdivision of the brain into two kinetically and anatomically distinct components, Brain1 and Brain2, plotted as in (A). (C) Area under the curve (AUC) for each component (injection site, CSF, Brain total, Brain1, and Brain2), compared between anesthesia groups. Boxplots show group distributions with individual pig values overlaid (point shapes as indicated in the legend: KD1–KD5, PRO1–PRO5). Asterisks indicate statistical significance from unpaired t-tests (* p < 0.05, ** p < 0.01). (D) Peak radioactivity concentration at the injection site and in the CSF component, compared between anesthesia groups (ns = not significant, unpaired t-test). (E) One-tissue compartment model (1TCM) schematic depicting CSF-to-brain tracer transport, parameterized by an influx rate constant (K_1_, CSF-to-brain, in µL⋅h⁻¹⋅g⁻¹) and an efflux rate constant (k_2_, brain-to-CSF, in h⁻¹). Boxplots show the estimated kinetic parameters K_1_, k_2_, and distribution volume (K_1_/k_2_=V_T_ in mL/cm^3^) for each anesthesia group, with individual pig values overlaid (ns = not significant, unpaired t-test).

Because the pig brain is gyrencephalic and contains many fissures and sulci, we investigated if the tracer distribution differed between brain areas. After decomposing the SPECT-data into 3 components (injection site, CSF and brain), we further subdivided the brain into more components using a hierarchical segmentation, obtaining two clear components (Figure 1D right) with different TAC patterns (Figure 2B). “Brain1” is spatially distributed in the border between the CSF and the brain especially in the ventral area and in the brain fissures and Brain2 is spatially confined in deeper areas of the brain and around the dorsal lateral hemispheres.

NMF-derived volumes of the CSF space and the brain parenchyma components were similar between the two groups: PRO (mean ± standard deviation (SD): 41.2 ± 5.1 cm^3^ for CSF and 68.8 ± 10.3 cm^3^ for brain) compared to the K/D group (37.8 ± 6.2 cm^3^ for CSF and 63.3± 8.4 cm^3^ for brain) [CSF and brain mean difference KD-PRO = −3.3 [−11.7, 5.0], and −5.5 [−19.3, 8.3] cm^3^ respectively; with p-values = 0.3 for both]. In both groups, the CSF and brain volumes represented ∼30% and ∼50%, respectively, of the total intracranial volume (sum of NMF components).

### 3.3 Intracranial tracer differences across anesthesia groups

Figure 2A shows TACs extracted from each NMF component, plotted individually per pig (by point shape) alongside anesthesia group means in Figure 2A, and with the hierarchical components in Figure 2B.

The AUC based on six hours was significantly larger for K/D than PRO at the injection site, cranial CSF space, the brain total and in the two brain sub-components, indicating higher tracer concentration in these regions over six hour period: estimated mean difference K/D-PRO [95% CI] injection site 0.38 [0.06, 0.7] normalized radioactivity concentration, p=0.026; CSF 0.18 [0.003, 0.4], p=0.047; brain total 0.06 [0.03, 0.09], p=0.002; Brain1 0.06 [0.02, 0.1], p=0.012; and Brain2 0.04 [0.01, 0.06], p=0.007. Those AUC differences were not driven by differences between groups in amount of tracer injected: K/D anesthetized pigs received a mean ± SD of 436 ± 54 MBq and PRO anesthetized pigs received 465 ± 43 MBq, p=0.3; and the groups had similar peak radioactivity from the injection site and CSF TACs (mean difference between K/D and PRO groups was -0.06 [-0.3, 0.2] normalized radioactivity concentration, p=0.6 and 0.02 [-0.02, 0.06], p=0.3, respectively) (Figure 2D). AUC and peak results comparison and effect sizes across anesthesia groups are shown in Table S3.

We found no statistically significant group differences in 1TCM kinetic parameters K_1_ (brain influx rate) or k_2_ (brain efflux rate). Mean plus SD of K_1_ for PRO was 98.92 ± 16.76 µL·cm^-3^·h^-1^ and for K/D was 118.71 ± 28.62 µL·cm^-3^·h^-1^; and k_2_ for PRO was 0.11 ± 0.06 h^-1^ and for K/D was 0.15 ± 0.06 h^-1^; with an estimate difference K/D-PRO in K_1_ of 19.78 [-14.43, 53.98] µL·cm^-3^·h^-1^, p=0.219 and k_2_ 0.03 [-0.06, 0.12] h⁻¹, p=0.481 (Figure 2E). KM comparison and effect sizes across anesthesia groups is included in Table S3. We also tested the suitability of a two-tissue compartment model (2TCM) model versus the 1TCM, but 2TCM did not improve confidence of parameter estimates nor model fit (results not shown), and accordingly, we chose the simplest model.

While using individual NMF maps may be more precise, it potentially biases a between-group analysis. As a sensitivity analysis, to assess any potential impact of individual anesthesia-related differences in the (3k)-NMF delineation, we also repeated the 1TCM using averaged component maps. We averaged the individual NMF maps using three different approaches: (1) averaging the individual NMF maps only from PRO pigs; (2) averaging the maps only from the K/D pigs; and (3) averaging across all pigs. Three-dimensional representations of these averaged component maps are shown in Supplementary Figure 1. Kinetic modelling using all three averaging approaches described above yielded the same conclusions as the individual component maps (Supplementary Figure 2). KM comparison and effects sizes across anesthesia groups for all NMF segmentations (individual and averages) are also included in Table S3.

### 3.4 Final tracer body distribution in the CNS across anesthesia groups

Final body radioactivity at the end of the brain scanning period showed no statistically significant anesthesia-specific tracer retention patterns between groups (Figure 3A). Despite differences in AUC from the CSF and brain TACs, final whole-body scans showed no difference across groups in cranial cavity tracer concentration. This discrepancy may reflect several factors: the lower sample size per group in this analysis (n = 4 vs. n = 5), differences in manual delineation between the cranial and spinal cord, or a dilution effect, whereby differences that accumulate across the full AUC become less detectable in a single timepoint measure. Spinal distribution, however, showed a trend toward greater tracer retention in the upper spinal cord with K/D than with PRO, in which the tracer spread more towards lower spinal regions, although this did not reach statistical significance. Moreover, since the whole-body scan was performed at the end of the experiment which for practical reasons could vary somewhat, we tested if timing differed between groups: estimate difference plus [95% CI] between K/D-PRO is -42 [-87.9, 3.9] minutes; p-value = 0.063. AUC from body scans comparison across anesthesia groups are included in Table S3.

**Figure 3.**
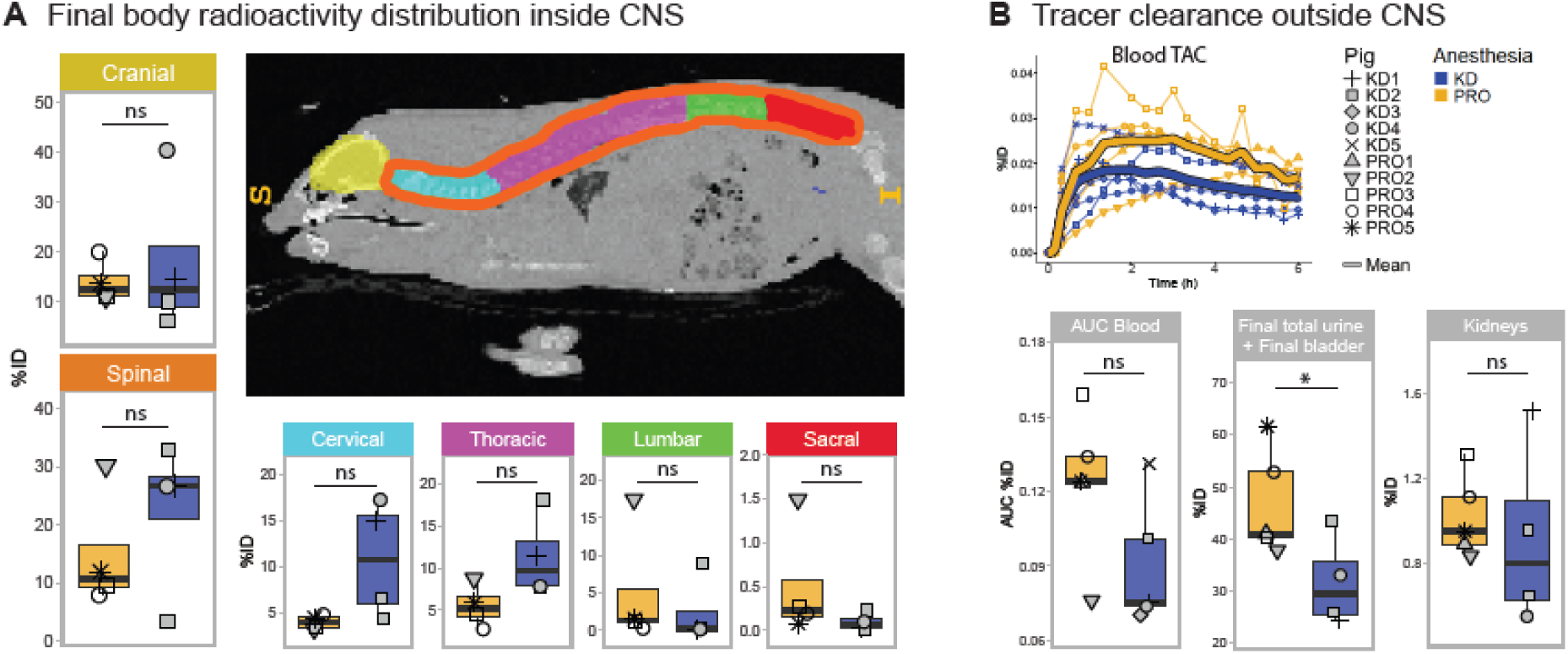
Whole-body tracer distribution within the CNS and systemic clearance under propofol and ketamine/dexmedetomidine anesthesia. A) Final whole-body radioactivity distribution within CNS regions of interest, expressed as percentage of injected dose (%ID) and compared between anesthesia groups: cranial, spinal cord (overall), and spinal cord subdivided into cervical, thoracic, lumbar, and sacral segments. Boxplots show group distributions with individual pig values overlaid (point shapes as indicated in the legend: KD1–KD5, PRO1–PRO5). Representative sagittal SPECT/CT image shows the manually delineated regions of interest by color (cranial, cervical, thoracic, lumbar, sacral) used for this analysis. (B) Tracer clearance outside the CNS. Top: blood time-activity curve (TAC, %ID) over the 6-hour scanning period, plotted per pig (thin lines with points) with group means overlaid (thick lines; blue = KD, orange = PRO). Bottom: area under the curve (AUC) for blood radioactivity, final total urine plus final bladder radioactivity (%ID), and kidney radioactivity (%ID), compared between anesthesia groups. Asterisks indicate statistical significance from unpaired t-tests (* p < 0.05, ns = not significant).

### 3.5 CNS tracer clearance across anesthesia groups

To assess tracer clearance from the CNS/CSF into the systemic distribution, we assessed whole blood radioactivity over time and the accumulated urine radioactivity collected during the whole experiment (Figure 3B). While K/D pigs generally had lower blood radioactivity, the only statistically significant difference we observed was that the PRO anesthetized pigs at the end of the experiment relative to the injected dose (%ID) had excreted significantly more radioactivity through the urine (40-60 %ID) than the K/D anesthetized pigs (25-40 %ID) [%ID estimate difference K/D-PRO = -22.1 [-38.0, -6.1], p=0.014]. CNS clearance results comparison across anesthesia groups are included in Table S3.

### 3.6 Kinetic modelling stability

When assessing the time stability of the kinetic modeling (KM) (Figure 4A), we found that k_2_ began to emerge between two- and five-hours post-injection and that with unconstrained TAC fitting, tracer brain efflux was slow, with a k_2_ of 0.13 ± 0.06 h^-1^ determined at six hours as an average across the ten pigs. K_1_ values were unstable with unconstrained data fitting (free k_2_), and since the estimation of K_1_ depends on the k_2_ estimation, to stabilize the K_1_ estimate, we tried fixing k_2_ values to either 0 h^-1^ or to 0.13 h^-1^. Fixing k_2_ to 0 did not decrease variability in K_1_ estimates, whereas fixing k_2_ to 0.13 improved the stability of K_1_ values from 2 h onwards (Figure 4A). Residual analysis of the 6 h kinetic modeling also demonstrated a better fit when fixing k_2_ at 0.13 h^-1^ compared with fixing k_2_ at 0 (Figure 4B). End-truncated KM pairwise comparison results (pooling anesthesia groups) for each k_2_ parameter are included in Table S4.

**Figure 4.**
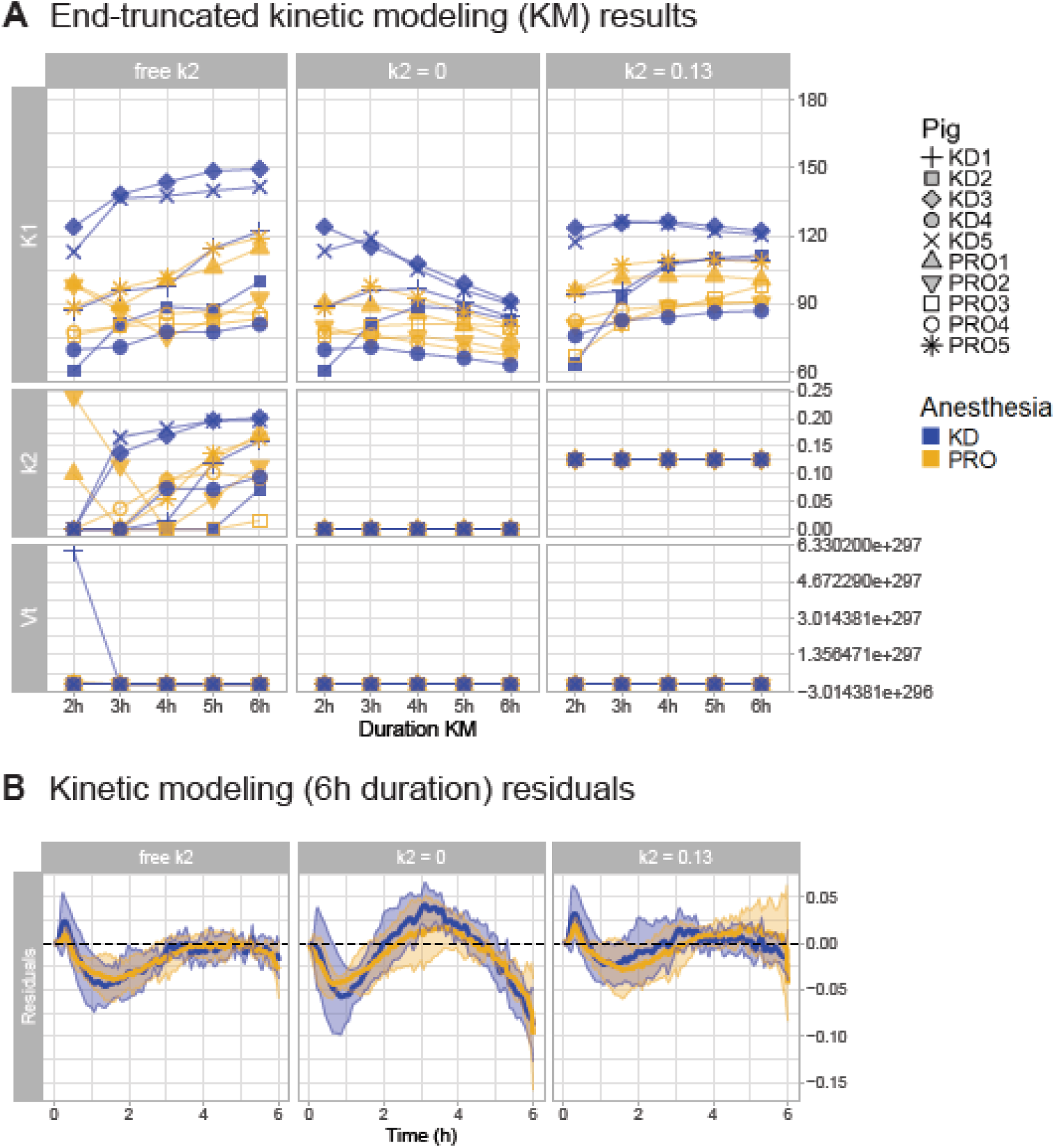
Stability of kinetic modelling for brain influx. A) End-truncation analysis showing kinetic parameters (K_1_ (µL⋅h⁻¹⋅cm⁻³), k_2_ (h^-1^), and V_T_ (mL/cm³)) stability with three modelling alternatives for the efflux parameter: k_2_ estimated freely, fixed at 0, or fixed at 0.13 h^-1^ (the mean k_2_ value at 6 hours post-injection), for each individual animal. B) Model residuals (in normalized radioactivity concentration units) for the three k_2_ alternatives described in A), derived from fits from the six-hour acquisition with mean plus SD for each group.

## 4. Discussion

In this study, we examine CSF and brain tracer distribution and kinetic differences in pigs undergoing two different types of anesthesia regimens: PRO, a GABAergic anesthetic; and ketamine/dexmedetomidine, an agent that rodent studies repeatedly have shown to increase tracer influx into the brain parenchyma^7,8,10–13^. We delineated CSF and brain spaces within the intracranial space using an automatic three-component NMF, and applied a 1TCM to characterize tracer movement into and out of the brain from the cranial CSF space.

For face validity, the two anesthetic regimens produced distinct EEG and physiological profiles in pigs that were consistent with their known pharmacological effects in humans^53–55^: Compared with PRO, K/D enhanced EEG-delta power and decreased alpha activity, resembling aspects of the EEG signature of deep non-REM sleep. DEX was also associated with higher mean arterial pressure and cerebral perfusion pressure, along with a lower heart rate, similar to what is seen in humans. Notably, no differences were observed in intracranial pressure or oxygen saturation between the two anesthetic conditions and the vitals remained relatively stable over the six hours. These findings support the translational relevance of the porcine model, indicating that the neurophysiological and systemic effects of these anesthetic agents are broadly comparable to those observed in humans.

Our study demonstrates that tracer injected into the cisterna magna accumulates more in brain tissue under K/D than under PRO anesthesia also in pigs, this difference was also detected in both subdivisions of the brain component. The observation aligns with prior rodent work, where K/D or ketamine plus another α_2_-adrenergic anesthetic, xylazine, are reported to lead to higher brain accumulation compared to isoflurane^4,7–13,21,56–61^ which, like propofol, is a GABAergic drug. But in contrast to a common interpretation that K/D promotes tracer influx to the brain parenchyma, in our study, the higher brain accumulation under K/D is not fully explained by an enhanced brain influx rate. Instead, it appears to be driven largely by a higher CSF tracer retention: basic kinetics predicts that a higher CSF tracer concentration increases proportionally to the amount of tracer entering the brain. Higher CSF tracer retention under K/D is not unique to our findings: Rodent studies have also reported higher AUC across the intracranial space (a signal dominated by CSF tracer) after intrathecal injection under K/D compared to isoflurane anesthesia^8,10^. This is also consistent with reported higher tracer retention in the cervical spinal cord under K/D^4,9^. The lower urinary tracer excretion that we observe with K/D compared to PRO anesthesia is also consistent with what has been reported in rodents: Lower tracer concentration in lymph nodes under K/D compared to isoflurane or the awake state^4,9,62^, and visually also a lower urinary excretion compared to isoflurane^8^. Although we do not find statistically significant differences in influx rates between anesthesia protocols, we cannot rule out that smaller, yet important, differences in K_1_ could contribute to a higher brain retention; such a demonstration would require a substantially higher sample size.

The higher CSF retention and lower peripheral excretion we observe in the K/D group are likely explained by a lower CSF turnover (defined as CSF production and drainage relative to CSF volume). Alternatively, since our K/D measurements are expressed relative to PRO, this could also be framed the other way: PRO may enhance CSF turnover. Correspondingly, higher CSF turnover under PRO would dilute the tracer within the CSF, drive it further down the spine, and increase peripheral excretion, consistent with what we observe. In rodents, direct measurement of CSF production (via cannulation of the lateral ventricle with occlusion of fourth-ventricular outflow) demonstrated 33% higher CSF production under isoflurane than under ketamine/xylazine anesthesia^63^. Although, direct data comparing CSF formation between K/D and PRO in gyrencephalic species are currently lacking, the porcine choroid plexus receive adrenergic innervation^64^, and porcine cerebral tissue expresses functional alpha2-adrenoceptors^65^. DEX-induced alpha2-adrenoceptor activation in the choroid plexus would be expected to reduce intracellular cAMP signaling, thereby decreasing the activity of ion transporters that drive water movement into the ventricles; α₂ receptor stimulation also suppresses Na⁺/K⁺-ATPase activity, which is central to the active ion transport that generates the osmotic gradient underlying CSF production.

With the NMF-based segmentation we did not find any anesthesia-related differences in brain or CSF volume that potentially could explain our observations, but we could be limited by the spatial resolution of a SPECT scanner. Also, simulations by Hornkjøl et al.^66^, revisited by Smets et al.^67^ suggest that while alterations in CSF turnover can explain our observations, changes in subarachnoid space volume alone cannot account for the anesthesia effects we observe in brain and CSF.

Methodologically, this study shows that combining (3k)-NMF segmentation with one-tissue compartment kinetic modelling offers a mechanistically informative framework for quantifying CSF tracer dynamics, providing extra information to the conventional AUC-based approaches for evaluating tracer influx to the brain. Further, the NMF approach was chosen to separate the CSF spaces from the brain tissue, which manually was not possible, and to reduce spillover/partial-volume contamination. Estimating tracer influx and efflux kinetics, allows for discrimination between distinct mechanisms that would otherwise be conflated into one measure.

Our hierarchical segmentation of the gyrencephalic pig brain provided a subdivision in two brain areas, Brain1 and Brain2, with Brain1 being located more superficially along the ventral brain surface, in the interhemispheric fissure and near the olfactory bulbs and cerebellum whereas Brain2 was comprised of deeper brain voxels plus surface voxels restricted to the dorsolateral hemispheres. This distribution is consistent with a study in mice demonstrating that infusion into the cisterna magna promotes transport toward ventral regions, whereas intraventricular injection favors transport across the ventricular-parenchymal interface, highlighting that tracer distribution is sensitive to injection place^13^. The two regions also closely match the tracer distribution pattern reported by Bèchet et al.^35^ in an *ex vivo* microscopy image of a pig brain, supporting the validity of our hierarchical segmentation.

Brain influx (K_1_) remained stable from 2 h post-injection onwards when fixing k_2_ to the average value estimated from 6 h dynamic modeling, and VT was very well determined. However, a limitation of our experimental setup is that pig brain activity peaks around 4 to 5 hours post-injection, meaning 6 hours of scanning is not sufficient to determine a stable efflux parameter (k_2_). This slow CSF dynamic differs greatly from rodent experiments, where 1 hour is sufficient to observe the radioactivity peak in the brain^11,68^. This species difference in tracer kinetics reflects the scaling of CSF volume and turnover rate with brain size: larger, gyrated brains require more time for tracer distribution and clearance, extending the timescale of all kinetic processes. In addition, the kinetic differences found between brain sub-divisions raise concerns about the interpretability of the estimated k_2_ parameter. Based on our data, it is not possible to determine whether the tracer diffuses directly from CSF to Brain2 or from surface areas (Brain1) to deeper regions (Brain2); and since the Brain2 TAC does not reach a plateau during the scanning period, we cannot be certain whether “k_2_” actually reflects brain efflux or tracer penetrating deeper from Brain1 into Brain2. To better assess tracer efflux from the brain in large mammals, longer scanning times or intraparenchymal tracer injection would be preferable.

We propose that our experimental platform can be used as an analytical framework for future intrathecal tracer studies in large animals and in humans. These findings have direct implications for the interpretation and planning of both preclinical rodent studies and clinical imaging protocols, for example, in the optimization of intrathecal drug delivery^12^, and offer potential future translational perspective into perioperative care and anesthesia-induced delirium, which has been associated with impaired brain clearance following prolonged anesthetic exposure^69–71^.

In summary, we here provide an experimental platform that can be used as an analytical framework for future intrathecal tracer studies in large animals and in humans. These findings have direct implications for the interpretation and planning of both preclinical rodent studies and clinical imaging protocols, for example, in the optimization of intrathecal drug delivery^12^, and offer potential future translational perspective into perioperative care and anesthesia-induced delirium, which has been associated with impaired brain clearance following prolonged anesthetic exposure^69–71^. Based on this platform, we show that the higher brain tracer accumulation with K/D compared to PRO is primarily explained by higher CSF tracer retention, likely resulting from a lower CSF turnover although we cannot rule out that brain influx rates to a lesser extent are also affected by anesthesia regime.

## Abbreviations

1TCM: One-tissue compartment model
AUC: Area under the curve
CM: Cisterna magna
CNS: Central nervous system
CPP: Cerebral perfusion pressure
CSF: Cerebrospinal fluid
CT: Computed tomography
DEX: Dexmedetomidine
EEG: Electroencephalogram
ICP: Intracranial pressure
K/D: Ketamine + dexmedetomidine
KM: Kinetic modelling
MAP: Mean arterial pressure
NMF: Non-negative matrix factorization
PRO: Propofol
ROI: Region of interest
SPECT: Single-photon emission computed tomography
TAC: Time activity curve
Tc^99m^-DTPA: Technetium-diethylenetriaminepentaacetic acid

---

**Supplementary Figure 1:**
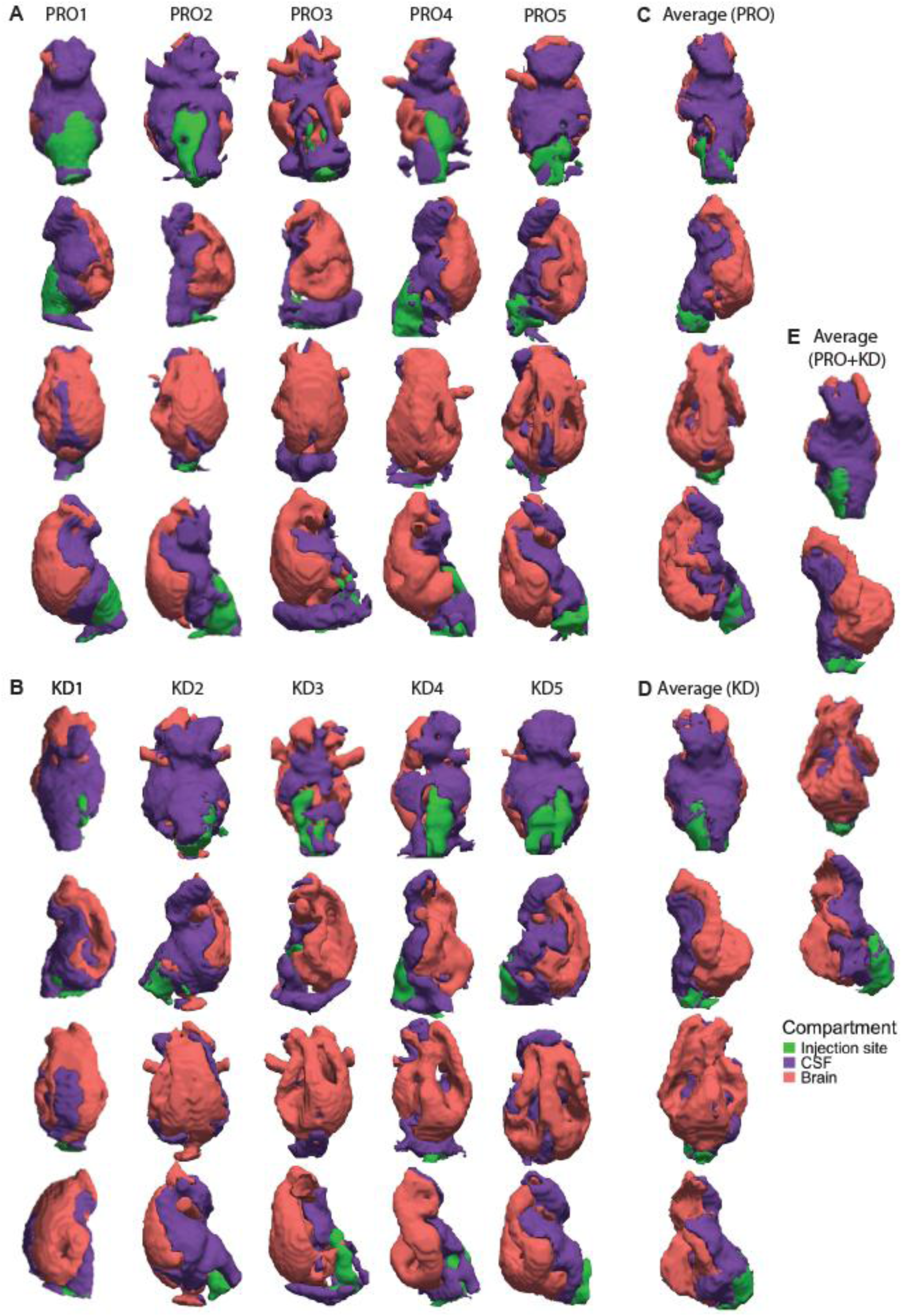
Individual and average 3D maps from (3k-)-NMF. A) Maps from PRO group (PRO1-5). B) Maps from K/D group (KD1-5). C) Average maps using only PRO maps. D) Average maps using only K/D maps. E) Average maps using all maps (PRO and K/D). Brain component in pink, CSF component in purple and injection site in green.

**Supplementary Figure 2:**
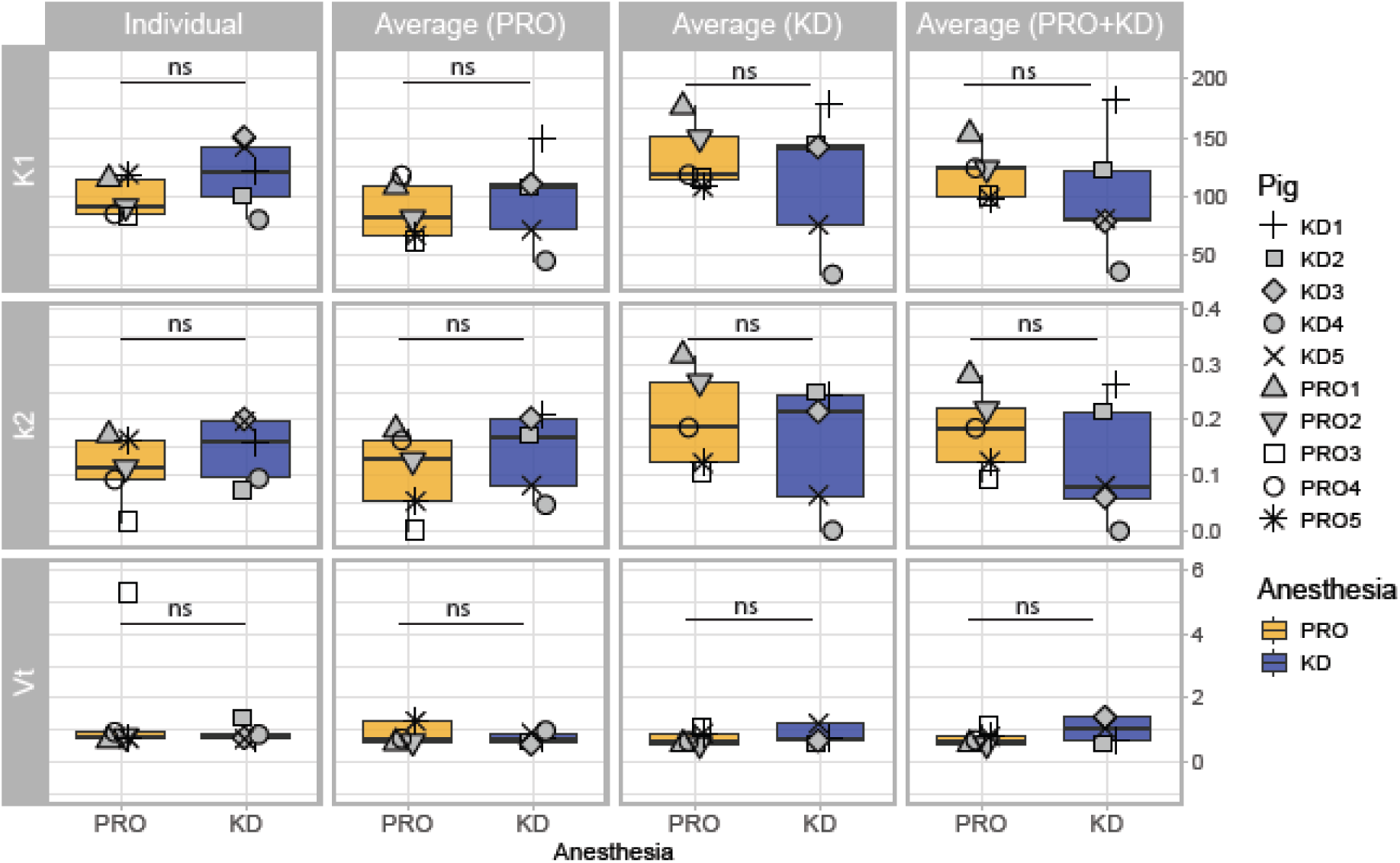
1TCM results for individual versus average component maps across pigs. Boxplots show the estimated kinetic parameters: influx rate constant (K_1_, CSF-to-brain, in µL⋅h⁻¹⋅g⁻¹) and an efflux rate constant (k_2_, brain-to-CSF, in h⁻¹), and distribution volume (V_T_, mL/cm^3^) for each anesthesia group, with individual pig values. None of the differences were statistically significant (unpaired t-test).

## References

1. Iliff JJ, Wang M, Liao Y, et al. A paravascular pathway facilitates CSF flow through the brain parenchyma and the clearance of interstitial solutes, including amyloid β. Sci Transl Med. 2012;4(147):147ra111. doi:10.1126/scitranslmed.3003748

2. Mestre H, Hablitz LM, Xavier AL, et al. Aquaporin-4-dependent glymphatic solute transport in the rodent brain. eLife. 2018;7. doi:10.7554/eLife.40070

3. Xie L, Kang H, Xu Q, et al. Sleep Drives Metabolite Clearance from the Adult Brain. Science. 2013;342(6156):373–377. doi:10.1126/science.1241224

4. Miyakoshi LM, Stæger FF, Li Q, et al. The state of brain activity modulates cerebrospinal fluid transport. Prog Neurobiol. 2023;229:102512. doi:10.1016/j.pneurobio.2023.102512

5. Gakuba C, Gaberel T, Goursaud S, et al. General Anesthesia Inhibits the Activity of the “Glymphatic System”. Theranostics. 2018;8(3):710–722. doi:10.7150/thno.19154

6. Lucey BP, Hicks TJ, McLeland JS, et al. Effect of sleep on overnight cerebrospinal fluid amyloid β kinetics. Ann Neurol. 2018;83(1):197–204. doi:10.1002/ana.25117

7. Hablitz LM, Vinitsky HS, Sun Q, et al. Increased glymphatic influx is correlated with high EEG delta power and low heart rate in mice under anesthesia. Sci Adv. 2019;5(2):eaav5447. doi:10.1126/sciadv.aav5447

8. Persson NDÅ, Lohela TJ, Mortensen KN, et al. Anesthesia Blunts Carbon Dioxide Effects on Glymphatic Cerebrospinal Fluid Dynamics in Mechanically Ventilated Rats. Anesthesiology. 2024;141(2):338–352. doi:10.1097/ALN.0000000000005039

9. Sun B, Fang D, Li W, Li M, Zhu S. NIR-II nanoprobes for investigating the glymphatic system function under anesthesia and stroke injury. J Nanobiotechnology. 2024;22(1):200. doi:10.1186/s12951-024-02481-w

10. Sigurdsson B, Hauglund NL, Lilius TO, et al. A SPECT-based method for dynamic imaging of the glymphatic system in rats. J Cereb Blood Flow Metab. 2023;43(7):1153–1165. doi:10.1177/0271678x231156982

11. Benveniste H, Lee H, Ding F, et al. Anesthesia with Dexmedetomidine and Low-dose Isoflurane Increases Solute Transport via the Glymphatic Pathway in Rat Brain When Compared with High-dose Isoflurane. Anesthesiology. 2017;127(6):976–988. doi:10.1097/ALN.0000000000001888

12. Lilius TO, Blomqvist K, Hauglund NL, et al. Dexmedetomidine enhances glymphatic brain delivery of intrathecally administered drugs. J Controlled Release. 2019;304:29–38. 10.1016/j.jconrel.2019.05.005

13. Zhu Y, Zhu J, Ni C, et al. Impact of infusion conditions and anesthesia on CSF tracer dynamics in mouse brain. Physiology. Preprint posted online January 24, 2025. doi:10.1101/2025.01.21.634133

14. Beschorner N, Navarro ML, Rosenholm M, et al. Scaling of anesthesia-dependent cerebrospinal fluid dynamics across rat and pig brains. Neuroscience. Preprint posted online July 16, 2026. doi:10.64898/2026.07.10.737206

15. Hauglund NL, Andersen M, Tokarska K, et al. Norepinephrine-mediated slow vasomotion drives glymphatic clearance during sleep. Cell. 2025;188(3):606–622.e17. doi:10.1016/j.cell.2024.11.027

16. Wang S, Yu X, Cheng L, et al. Dexmedetomidine improves the circulatory dysfunction of the glymphatic system induced by sevoflurane through the PI3K/AKT/ΔFosB/AQP4 pathway in young mice. Cell Death Dis. 2024;15(6):448. doi:10.1038/s41419-024-06845-w

17. Li J, Liu X, Bo B, et al. Glymphatic influx is negatively correlated with cerebral blood volume in male mice. Cell Rep. 2026;45(4):117182. doi:10.1016/j.celrep.2026.117182

18. Zhang R. A Comprehensive Review of the Depth of Anesthesia and Its Impact on Postoperative Delirium Development. Theor Nat Sci. 2025;85(1):172–177. doi:10.54254/2753-8818/2025.20186

19. Inouye SK. Delirium in Older Persons. N Engl J Med. 2006;354(11):1157–1165. doi:10.1056/NEJMra052321

20. Saczynski JS, Inouye SK, Kosar CM, et al. Cognitive and brain reserve and the risk of postoperative delirium in older patients: analysis of data from a prospective observational study. Lancet Psychiatry. 2014;1(6):437–443. doi:10.1016/S2215-0366(14)00009-1

21. Sun X, Fang D, Zhang X, et al. Engineering Near-Infrared-II Nanoprobes Reveal Dexmedetomidine Potentiating Brain Waste Clearance in Healthy and Sleep-Restricted Mice. ACS Nano. 2025;19(39):34830–34846. doi:10.1021/acsnano.5c10519

22. MacLaren R, Preslaski CR, Mueller SW, et al. A Randomized, Double-Blind Pilot Study of Dexmedetomidine Versus Midazolam for Intensive Care Unit Sedation: Patient Recall of Their Experiences and Short-Term Psychological Outcomes. J Intensive Care Med. 2015;30(3):167–175. doi:10.1177/0885066613510874

23. Duan X, Coburn M, Rossaint R, Sanders RD, Waesberghe JV, Kowark A. Efficacy of perioperative dexmedetomidine on postoperative delirium: systematic review and meta-analysis with trial sequential analysis of randomised controlled trials. Br J Anaesth. 2018;121(2):384–397. doi:10.1016/j.bja.2018.04.046

24. Liu Y, Li XJ, Liang Y, Kang Y. Pharmacological Prevention of Postoperative Delirium: A Systematic Review and Meta-Analysis of Randomized Controlled Trials. Evid Based Complement Alternat Med. 2019;2019:1–10. doi:10.1155/2019/9607129

25. Fultz NE, Bonmassar G, Setsompop K, et al. Coupled electrophysiological, hemodynamic, and cerebrospinal fluid oscillations in human sleep. Science. 2019;366(6465):628–631. doi:10.1126/science.aax5440

26. Vijayakrishnan Nair V, Kish BR, Inglis B, et al. Human CSF movement influenced by vascular low frequency oscillations and respiration. Front Physiol. 2022;13:940140. doi:10.3389/fphys.2022.940140

27. Dreha-Kulaczewski S, Joseph AA, Merboldt KD, Ludwig HC, Gärtner J, Frahm J. Inspiration Is the Major Regulator of Human CSF Flow. J Neurosci. 2015;35(6):2485–2491. doi:10.1523/JNEUROSCI.3246-14.2015

28. Eide PK, Vinje V, Pripp AH, Mardal KA, Ringstad G. Sleep deprivation impairs molecular clearance from the human brain. Brain. 2021;144(3):863–874. doi:10.1093/brain/awaa443

29. Vinje V, Zapf B, Ringstad G, Eide PK, Rognes ME, Mardal KA. Human brain solute transport quantified by glymphatic MRI-informed biophysics during sleep and sleep deprivation. Fluids Barriers CNS. 2023;20(1):62. doi:10.1186/s12987-023-00459-8

30. Valnes LM, Mitusch SK, Ringstad G, Eide PK, Funke SW, Mardal KA. Apparent diffusion coefficient estimates based on 24 hours tracer movement support glymphatic transport in human cerebral cortex. Sci Rep. 2020;10(1):9176. doi:10.1038/s41598-020-66042-5

31. Howells DW, Porritt MJ, Rewell SSJ, et al. Different strokes for different folks: the rich diversity of animal models of focal cerebral ischemia. J Cereb Blood Flow Metab Off J Int Soc Cereb Blood Flow Metab. 2010;30(8):1412–1431. doi:10.1038/jcbfm.2010.66

32. Kobayashi E, Hishikawa S, Teratani T, Lefor AT. The pig as a model for translational research: overview of porcine animal models at Jichi Medical University. Transplant Res. 2012;1(1):8. doi:10.1186/2047-1440-1-8

33. Lunney JK, Van Goor A, Walker KE, Hailstock T, Franklin J, Dai C. Importance of the pig as a human biomedical model. Sci Transl Med. 2021;13(621):eabd5758.

34. Robert S, Dallaire A. Polygraphic analysis of the sleep-wake states and the REM sleep periodicity in domesticated pigs (Sus scrofa). Physiol Behav. 1986;37(2):289–293. doi:10.1016/0031-9384(86)90235-0

35. Bèchet NB, Shanbhag NC, Lundgaard I. Glymphatic pathways in the gyrencephalic brain. J Cereb Blood Flow Metab Off J Int Soc Cereb Blood Flow Metab. 2021;41(9):2264–2279. doi:10.1177/0271678X21996175

36. Donovan LL, Johansen JV, Ros NF, et al. Effects of a single dose of psilocybin on behaviour, brain 5-HT(2A) receptor occupancy and gene expression in the pig. Eur Neuropsychopharmacol J Eur Coll Neuropsychopharmacol. 2021;42:1–11. doi:10.1016/j.euroneuro.2020.11.013

37. Raval NR, Nasser A, Madsen CA, et al. An in vivo Pig Model for Testing Novel Positron Emission Tomography Radioligands Targeting Cerebral Protein Aggregates. Front Neurosci. 2022;16:847074. doi:10.3389/fnins.2022.847074

38. Villadsen J, Hansen HD, Jørgensen LM, et al. Automatic delineation of brain regions on MRI and PET images from the pig. J Neurosci Methods. 2018;294:51–58. doi:10.1016/j.jneumeth.2017.11.008

39. Jørgensen LM, Weikop P, Villadsen J, et al. Cerebral 5-HT release correlates with [(11)C]Cimbi36 PET measures of 5-HT2A receptor occupancy in the pig brain. J Cereb Blood Flow Metab Off J Int Soc Cereb Blood Flow Metab. 2017;37(2):425–434. doi:10.1177/0271678X16629483

40. Sauleau P, Lapouble E, Val-Laillet D, Malbert CH. The pig model in brain imaging and neurosurgery. Anim Int J Anim Biosci. 2009;3(8):1138–1151. doi:10.1017/S1751731109004649

41. Andersen OM, Bøgh N, Landau AM, et al. A genetically modified minipig model for Alzheimer’s disease with SORL1 haploinsufficiency. Cell Rep Med. 2022;3(9):100740. doi:10.1016/j.xcrm.2022.100740

42. Percie du Sert N, Hurst V, Ahluwalia A, et al. The ARRIVE guidelines 2.0: Updated guidelines for reporting animal research. PLoS Biol. 2020;18(7):e3000410. doi:10.1371/journal.pbio.3000410

43. Zavorsky GS, Van Wijk XMR. The stability of blood gases and CO-oximetry under slushed ice and room temperature conditions. Clin Chem Lab Med CCLM. 2023;61(10):1750–1759. doi:10.1515/cclm-2022-1085

44. Thakur V, Akerele OA, Randell E. Validation of glucose and lactate in cerebrospinal fluid (CSF) on a Radiometer blood gas analyzer ABL90 Flex plus. Clin Biochem. 2025;136:110876. doi:10.1016/j.clinbiochem.2025.110876

45. Lee HC, Jung CW. Vital Recorder—a free research tool for automatic recording of high-resolution time-synchronised physiological data from multiple anaesthesia devices. Sci Rep. 2018;8(1):1527. doi:10.1038/s41598-018-20062-4

46. Prerau MJ, Brown RE, Bianchi MT, Ellenbogen JM, Purdon PL. Sleep Neurophysiological Dynamics Through the Lens of Multitaper Spectral Analysis. Physiology. 2017;32(1):60–92. doi:10.1152/physiol.00062.2015

47. Lee DD, Seung HS. Learning the parts of objects by non-negative matrix factorization. Nature. 1999;401(6755):788–791. doi:10.1038/44565

48. Olsen AS, Navarro ML, Svarer C, Hinrich JL, Mørup M, Knudsen GM. Shift-and Stretch-Invariant Non-Negative Matrix Factorization with an Application to Brain Tissue Delineation in Emission Tomography Data. In: ICASSP 2026 - 2026 IEEE International Conference on Acoustics, Speech and Signal Processing (ICASSP). IEEE; 2026:336-340. doi:10.1109/ICASSP55912.2026.11461301

49. Yushkevich PA, Piven J, Hazlett HC, et al. User-guided 3D active contour segmentation of anatomical structures: Significantly improved efficiency and reliability. NeuroImage. 2006;31(3):1116–1128. doi:10.1016/j.neuroimage.2006.01.015

50. Borchers HW. pracma: Practical Numerical Math Functions. Published online March 19, 2011:2.4.6. doi:10.32614/CRAN.package.pracma

51. OpenAI. ChatGPT. 2024. https://chat.openai.com

52. Claude. Claude. Accessed May 26, 2026. https://claude.ai/new

53. Reel B, Maani C. Dexmedetomidine. Treasure Island (FL): StatPearls Publishing; 2026. https://www.ncbi.nlm.nih.gov/books/NBK513303/

54. Weerink MAS, Struys MMRF, Hannivoort LN, Barends CRM, Absalom AR, Colin P. Clinical Pharmacokinetics and Pharmacodynamics of Dexmedetomidine. Clin Pharmacokinet. 2017;56(8):893–913. doi:10.1007/s40262-017-0507-7

55. Ebert TJ, Hall JE, Barney JA, Uhrich TD, Colinco MD. The Effects of Increasing Plasma Concentrations of Dexmedetomidine in Humans. Anesthesiology. 2000;93(2):382–394. doi:10.1097/00000542-200008000-00016

56. Choi S, Kim J, Jeon H, et al. Photoacoustic computed tomography monitors cerebrospinal fluid dynamics and glymphatic function. Nat Commun. 2026;17(1):2677. doi:10.1038/s41467-026-69390-4

57. Ge H, Gu X, Wang Z, et al. Anesthetics Modulate Cerebrospinal Fluid Efflux Pathways in Mice by Altering Perineural and Perivascular Spaces. NMR Biomed. 2026;39(2):e70222. doi:10.1002/nbm.70222

58. Segeroth M, Wachsmuth L, Gagel M, Albers F, Hess A, Faber C. Disentangling the impact of cerebrospinal fluid formation and neuronal activity on solute clearance from the brain. Fluids Barriers CNS. 2023;20(1):43. doi:10.1186/s12987-023-00443-2

59. Stanton EH, Persson NDÅ, Gomolka RS, et al. Mapping of CSF transport using high spatiotemporal resolution dynamic contrast-enhanced MRI in mice: Effect of anesthesia. Magn Reson Med. 2021;85(6):3326–3342. doi:10.1002/mrm.28645

60. Persson NDÅ, Lohela TJ, Anttila JE, et al. General Anesthesia and Systemic Hyperosmolality Modulate Lumbar Intrathecal Drug Distribution in Female Rats. Anesthesiology. 2026;144(2):390–401. doi:10.1097/ALN.0000000000005794

61. Tong T, Newbold E, Wang J, et al. Enhanced glymphatic CSF tracer influx during á2-adrenergic agonist anesthesia is independent of tracer injection duration. Neuroscience. Preprint posted online June 2, 2026. doi:10.64898/2026.05.29.728816

62. Ma Q, Ries M, Decker Y, et al. Rapid lymphatic efflux limits cerebrospinal fluid flow to the brain. Acta Neuropathol (Berl). 2019;137(1):151–165. doi:10.1007/s00401-018-1916-x

63. Liu G, Mestre H, Sweeney AM, et al. Direct Measurement of Cerebrospinal Fluid Production in Mice. Cell Rep. 2020;33(12):108524. doi:10.1016/j.celrep.2020.108524

64. Nilsson C, Kannisto P, Lindvall-Axelsson M, Owman C, Rosengren E. The neuropeptides vasoactive intestinal polypeptide, peptide histidine isoleucine and neuropeptide Y modulate [3H]noradrenaline release from sympathetic nerves in the choroid plexus. Eur J Pharmacol. 1990;181(3):247–252. doi:10.1016/0014-2999(90)90085-K

65. Asada Y, Lee TJ. Alpha 2-adrenoceptors mediate norepinephrine constriction of porcine pial veins. Am J Physiol-Heart Circ Physiol. 1992;263(6):H1907–H1910. doi:10.1152/ajpheart.1992.263.6.H1907

66. Hornkjøl M, Valnes LM, Ringstad G, et al. CSF circulation and dispersion yield rapid clearance from intracranial compartments. Front Bioeng Biotechnol. 2022;10:932469. doi:10.3389/fbioe.2022.932469

67. Smets NG, Strijkers GJ, Vinje V, Bakker ENTP. Cerebrospinal fluid turnover as a driver of brain clearance. NMR Biomed. 2024;37(7):e5029. doi:10.1002/nbm.5029

68. Mortensen KN, Sanggaard S, Mestre H, et al. Impaired Glymphatic Transport in Spontaneously Hypertensive Rats. J Neurosci. 2019;39(32):6365–6377. doi:10.1523/JNEUROSCI.1974-18.2019

69. Maldonado JR, Wysong A, Van Der Starre PJA, Block T, Miller C, Reitz BA. Dexmedetomidine and the Reduction of Postoperative Delirium after Cardiac Surgery. Psychosomatics. 2009;50(3):206–217. doi:10.1176/appi.psy.50.3.206

70. Lin Y, Chen J, Wang Z. Meta-Analysis of Factors Which Influence Delirium Following Cardiac Surgery. J Card Surg. 2012;27(4):481–492. doi:10.1111/j.1540-8191.2012.01472.x

71. Pandharipande PP, Pun BT, Herr DL, et al. Effect of Sedation With Dexmedetomidine vs Lorazepam on Acute Brain Dysfunction in Mechanically Ventilated Patients: The MENDS Randomized Controlled Trial. JAMA. 2007;298(22):2644. doi:10.1001/jama.298.22.2644

