## Supplementary Tables for "Kinetic analysis of CSF–brain tracer exchange in the pig brain under different anesthetic regimes"

### Table S1. Physiological vitals across pigs and groups

| Pig # | Group | MAP (*) | ICP | CPP (***) | ART. Diastolic (**) | ART. Systolic (*) | Pulse Pressure | Heart Rate (*) | $O_2$ saturation | $EtCO_2$ | Resp. volume | Resp. frequency | Blood Glucose |
|---|---|---|---|---|---|---|---|---|---|---|---|---|---|
| PRO1 | PRO | 57.2 ± 6.9 | - | - | 42.4 ± 8.0 | 90.0 ± 13.0 | 47.6 ± 12.8 | 79.9 ± 8.7 | 94.6 ± 2.1 | 5.8 ± 0.3 | 200.0 ± 0.0 | 14.9 ± 0.8 | 6.0 ± 3.4 |
| PRO2 | PRO | 82.2 ± 10.6 | 23.5 ± 2.6 | 58.8 ± 10.7 | 59.6 ± 4.3 | 117.6 ± 6.6 | 58.0 ± 4.7 | 94.4 ± 10.2 | 99.3 ± 1.4 | 5.6 ± 0.2 | 250.0 ± 0.0 | 20.0 ± 0.0 | 5.2 ± 0.6 |
| PRO3 | PRO | 65.0 ± 9.3 | 7.3 ± 2.4 | 57.8 ± 9.6 | 43.7 ± 1.8 | 100.6 ± 1.1 | 56.9 ± 1.6 | 78.6 ± 3.5 | 99.9 ± 0.6 | 6.0 ± 0.2 | 200.0 ± 0.0 | 20.5 ± 1.0 | 4.7 ± 0.6 |
| PRO4 | PRO | 73.0 ± 5.2 | 6.1 ± 0.6 | 66.9 ± 5.2 | 57.3 ± 4.2 | 97.3 ± 30.6 | 40.0 ± 31.3 | 107.3 ± 3.9 | 96.5 ± 0.5 | 6.1 ± 0.1 | 270.0 ± 0.0 | 22.0 ± 0.0 | 5.3 ± 0.7 |
| PRO5 | PRO | 72.3 ± 26.7 | 6.3 ± 1.2 | 65.5 ± 27.6 | 48.9 ± 3.6 | 99.9 ± 2.4 | 51.0 ± 2.0 | 80.0 ± 9.5 | 98.1 ± 0.7 | 5.4 ± 0.4 | 201.0 ± 8.8 | 18.6 ± 0.8 | 5.8 ± 2.4 |
| GROUP MEAN | PRO | 69.9 ± 9.4↓ | 10.8 ± 8.5 | 62.3 ± 4.6 | 50.4 ± 7.8↓ | 101.1 ± 10.2 | 50.7 ± 7.4 | 88.0 ± 12.6 | 97.7 ± 2.1 | 5.8 ± 0.3 | 224.2 ± 33.4 | 19.2 ± 2.7 | 5.4 ± 0.5 |
| KD1 | KD | 75.4 ± 8.5 | - | - | 63.1 ± 11.9 | 95.8 ± 7.9 | 34.6 ± 3.3 | 65.4 ± 6.9 | 98.7 ± 1.7 | 5.6 ± 0.3 | 142.9 ± 13.1 | 18.3 ± 0.9 | 5.7 ± 1.3 |
| KD2 | KD | 98.2 ± 12.9 | 10.6 ± 2.3 | 87.7 ± 12.5 | 77.9 ± 10.5 | 130.8 ± 10.2 | 52.9 ± 3.5 | 84.2 ± 9.0 | 96.9 ± 1.7 | 5.7 ± 0.2 | 232.0 ± 15.2 | 17.3 ± 0.7 | 9.8 ± 1.7 |
| KD3 | KD | 117.0 ± 14.3 | 17.4 ± 15.3 | 100.4 ± 17.7 | 95.8 ± 5.3 | 145.0 ± 8.3 | 49.2 ± 3.7 | 67.0 ± 5.8 | 99.6 ± 0.7 | 5.3 ± 0.4 | 231.9 ± 6.0 | 20.8 ± 5.7 | 5.6 ± 2.2 |
| KD4 | KD | 107.8 ± 2.7 | 5.4 ± 0.6 | 102.2 ± 1.5 | 90.3 ± 3.2 | 134.5 ± 3.7 | 44.3 ± 1.6 | 50.4 ± 2.5 | 98.7 ± 1.4 | 5.8 ± 0.3 | 215.0 ± 7.7 | 16.3 ± 0.5 | 9.6 ± 3.7 |
| KD5 | KD | 95.5 ± 5.8 | 3.1 ± 2.1 | 92.6 ± 7.1 | 73.5 ± 4.9 | 135.2 ± 9.4 | 61.7 ± 5.3 | 77.7 ± 25.3 | 99.9 ± 0.2 | 5.6 ± 0.3 | 200.0 ± 18.5 | 19.6 ± 0.7 | 9.1 ± 6.5 |
| GROUP MEAN | KD | 98.8 ± 15.6 | 9.1 ± 6.3 | 95.7 ± 6.8 | 80.1 ± 13.1 | 128.2 ± 18.9↑ | 48.5 ± 10.0 | 68.9 ± 12.9↓ | 98.8 ± 1.2 | 5.6 ± 0.2 | 204.4 ± 36.9↑ | 18.5 ± 1.8 | 7.9 ± 2.1 |

Values represent mean ± standard deviation (SD) across the 6-hour scanning period for each physiological parameter, shown per pig and as group means. Pulse pressure is calculated as the difference between systolic and diastolic arterial pressure. Cerebral perfusion pressure (CPP) is calculated as the difference between MAP and ICP. Units are as follows: mean arterial pressure (MAP), intracranial pressure (ICP), cerebral perfusion pressure (CPP), diastolic pressure (ART. Diastolic), systolic pressure (ART. Systolic), and pulse pressure in mmHg; heart rate in beats per minute; $O_2$ saturation in %; EtCO2 in kPa; respiration volume in mL; respiration frequency in breaths per minute; and blood glucose in mmol/L. Column headers indicate the statistical significance of an unpaired t-test comparing group means ('empty' = not significant; * $p < 0.05$; ** $p < 0.01$; *** $p < 0.001$). Within group mean rows, up and down arrows indicate whether each parameter increased or decreased, respectively, from the first half of scanning (0–3 h) to the second half (3–6 h); the number of arrows reflects the significance level of the corresponding paired t-test ('empty' = not significant; single arrow = $p < 0.05$, double = $p < 0.01$, triple = $p < 0.001$).

**Table S2. EEG bandwidth across pigs and groups**

| Pig # | Group | Alpha (***) | Beta (***) | Delta (***) | Gamma | Theta (**) |
|---|---|---|---|---|---|---|
| PRO1 | PRO | 0.094 ± 0.009 | 0.073 ± 0.012 | 0.518 ± 0.031 | 0.017 ± 0.006 | 0.265 ± 0.013 |
| PRO2 | PRO | 0.137 ± 0.007 | 0.108 ± 0.008 | 0.444 ± 0.020 | 0.035 ± 0.007 | 0.247 ± 0.012 |
| PRO3 | PRO | 0.112 ± 0.005 | 0.075 ± 0.008 | 0.522 ± 0.029 | 0.022 ± 0.006 | 0.239 ± 0.024 |
| PRO4 | PRO | 0.094 ± 0.023 | 0.078 ± 0.026 | 0.565 ± 0.073 | 0.009 ± 0.001 | 0.217 ± 0.039 |
| PRO5 | PRO | 0.099 ± 0.009 | 0.079 ± 0.004 | 0.527 ± 0.027 | 0.010 ± 0.002 | 0.260 ± 0.024 |
| GROUP MEAN | PRO | 0.107 ± 0.018 | 0.083 ± 0.014 | 0.515 ± 0.044 | 0.018 ± 0.010 | 0.246 ± 0.019 |
| KD1 | KD | 0.049 ± 0.008 | 0.017 ± 0.003 | 0.659 ± 0.021 | 0.004 ± 0.004 | 0.173 ± 0.024 |
| KD2 | KD | 0.036 ± 0.014 | 0.012 ± 0.005 | 0.717 ± 0.044 | 0.007 ± 0.001 | 0.121 ± 0.015 |
| KD3 | KD | 0.043 ± 0.011 | 0.016 ± 0.002 | 0.699 ± 0.032 | 0.005 ± 0.000 | 0.114 ± 0.013 |
| KD4 | KD | 0.071 ± 0.007 | 0.027 ± 0.001 | 0.632 ± 0.035 | 0.008 ± 0.001 | 0.206 ± 0.030 |
| KD5 | KD | 0.051 ± 0.006 | 0.020 ± 0.003 | 0.675 ± 0.022 | 0.005 ± 0.001 | 0.180 ± 0.029 |
| GROUP MEAN | KD | 0.050 ± 0.013 | 0.018 ± 0.006 | 0.676 ± 0.033 | 0.006 ± 0.001 | 0.159 ± 0.040 |

Values represent mean ± standard deviation (SD) across the 6-hour scanning period for each EEG frequency band, shown per pig and as group means. Relative band power is reported for the alpha, beta, gamma, delta, and theta frequency ranges, each expressed as a relative proportion of the total power. Column headers indicate the statistical significance of an unpaired t-test comparing group means (‘empty’ = not significant; * $p < 0.05$; ** $p < 0.01$; *** $p < 0.001$). Within group mean rows, up and down arrows indicate whether each band's relative power increased or decreased, respectively, from the first half of scanning (0–3 h) to the second half (3–6 h); the number of arrows reflects the significance level of the corresponding paired t-test (‘empty’ = not significant; single arrow = $p < 0.05$, double = $p < 0.01$, triple = $p < 0.001$).

**Table S3. Anesthesia comparison results (intracranial NMF AUC, TAC peak, body AUC, and KM for the different NMF segmentations)**

| Data | Analysis | Unit | ROI/Parameter | N (KD) | N (PRO) | Mean (KD) | Mean (PRO) | Mean difference (K/D-PRO) [95% CI] | T-statistic | DF | p.value | Signif. | Cohens' d [95% CI] |
|---|---|---|---|---|---|---|---|---|---|---|---|---|---|
| NMF TAC | AUC | Norm. [Radioct.] | Injection site | 5 | 5 | 1.258 | 0.881 | 0.38 [0.06, 0.69] | 2.87 | 7 | 0.026 | * | 1.8 [0.1, 3.5] |
| NMF TAC | AUC | Norm. [Radioct.] | CSF | 5 | 5 | 0.533 | 0.350 | 0.18 [0, 0.36] | 2.36 | 8 | 0.047 | * | 1.5 [-0.2, 3.1] |
| NMF TAC | AUC | Norm. [Radioct.] | Brain | 5 | 5 | 0.162 | 0.100 | 0.06 [0.03, 0.09] | 4.37 | 8 | 0.003 | ** | 2.8 [0.7, 4.8] |
| NMF TAC | AUC | Norm. [Radioct.] | Brain1 | 5 | 5 | 0.177 | 0.116 | 0.06 [0.02, 0.1] | 3.36 | 7 | 0.012 | * | 2.1 [0.3, 3.9] |
| NMF TAC | AUC | Norm. [Radioct.] | Brain2 | 5 | 5 | 0.099 | 0.063 | 0.04 [0.01, 0.06] | 3.63 | 8 | 0.007 | ** | 2.3 [0.4, 4.2] |
| NMF TAC | Peak | Norm. [Radioct.] | Injection site | 5 | 5 | 0.758 | 0.814 | -0.06 [-0.33, 0.22] | -0.55 | 4 | 0.611 | ns | -0.3 [-1.8, 1.1] |
| NMF TAC | Peak | Norm. [Radioct.] | CSF | 5 | 5 | 0.120 | 0.102 | 0.02 [-0.02, 0.06] | 1.06 | 7 | 0.327 | ns | 0.7 [-0.8, 2.2] |
| NMF TAC | Peak | Norm. [Radioct.] | Brain | 5 | 5 | 0.037 | 0.023 | 0.01 [0.01, 0.02] | 3.73 | 8 | 0.006 | ** | 2.4 [0.5, 4.3] |
| NMF TAC | Peak | Norm. [Radioct.] | Brain1 | 5 | 5 | 0.042 | 0.027 | 0.02 [0.01, 0.02] | 4.70 | 7 | 0.002 | ** | 3 [0.9, 5.1] |
| NMF TAC | Peak | Norm. [Radioct.] | Brain2 | 5 | 5 | 0.031 | 0.020 | 0.01 [0, 0.02] | 3.61 | 7 | 0.008 | ** | 2.3 [0.4, 4.2] |
| Body scan TAC | AUC | %ID | Cranial | 4 | 4 | 18.57 | 15.61 | 2.97 [-16.06, 21.99] | 0.45 | 4 | 0.676 | ns | 0.3 [-1.4, 2.1] |
| Body scan TAC | AUC | %ID | Spinal | 4 | 4 | 26.26 | 16.70 | 9.56 [-16.08, 35.2] | 0.95 | 5 | 0.384 | ns | 0.7 [-1.1, 2.5] |
| Body scan TAC | AUC | %ID | Kidneys | 4 | 5 | 1.097 | 1.291 | -0.19 [-1.16, 0.77] | -0.53 | 5 | 0.622 | ns | -0.4 [-2, 1.2] |
| Body scan TAC + Urine | AUC | %ID | Urine+Bladder | 4 | 5 | 35.78 | 57.86 | -22.08 [-38.02, -6.13] | -3.27 | 7 | 0.014 | * | -2.1 [-4.1, -0.1] |
| Blood ex vivo | AUC | %ID | Blood | 5 | 5 | 0.090 | 0.123 | -0.03 [-0.07, 0.01] | -1.86 | 8 | 0.101 | ns | -1.2 [-2.8, 0.4] |
| Body scan TAC | AUC | %ID | Cervical | 4 | 4 | 12.01 | 4.55 | 7.46 [-3.08, 17.99] | 2.23 | 3 | 0.111 | ns | 1.6 [-0.4, 3.6] |
| Body scan TAC | AUC | %ID | Thoracic | 4 | 4 | 13.57 | 6.21 | 7.36 [-4.02, 18.74] | 1.84 | 4 | 0.144 | ns | 1.3 [-0.6, 3.2] |
| Body scan TAC | AUC | %ID | Lumbar | 4 | 4 | 3.085 | 5.611 | -2.53 [-15.99, 10.94] | -0.48 | 5 | 0.652 | ns | -0.3 [-2.1, 1.4] |
| Body scan TAC | AUC | %ID | Sacral | 4 | 4 | 0.113 | 0.569 | -0.46 [-1.56, 0.65] | -1.27 | 3 | 0.289 | ns | -0.9 [-2.7, 0.9] |
| Body scan | minutes | minutes | Scan time | 4 | 4 | 445.5 | 487.5 | -42 [-87.94, 3.94] | -2.67 | 4 | 0.063 | . | -1.9 [-4, 0.2] |
| Individual NMF | KM | $\mu L \cdot cm^{-3} \cdot h^{-1}$ | K1 | 5 | 5 | 118.71 | 98.93 | 19.78 [-14.43, 53.98] | 1.3 | 8 | 0.219 | ns | 0.84 [-0.69, 2.38] |
| Individual NMF | KM | $h^{-1}$ | k2 | 5 | 5 | 0.145 | 0.112 | 0.03 [-0.06, 0.12] | 0.9 | 8 | 0.416 | ns | 0.54 [-0.95, 2.03] |
| Individual NMF | KM | $mL/cm^3$ | Vt | 5 | 5 | 0.891 | 1.682 | -0.79 [-2.89, 1.31] | -0.9 | 8 | 0.411 | ns | -0.55 [-2.04, 0.94] |
| Average (PRO pigs) | KM | $\mu L \cdot cm^{-3} \cdot h^{-1}$ | K1 | 5 | 5 | 97.24 | 87.76 | 9.49 [-38.92, 57.89] | 0.5 | 8 | 0.663 | ns | 0.29 [-1.18, 1.75] |
| Average (PRO pigs) | KM | $h^{-1}$ | k2 | 5 | 5 | 0.142 | 0.105 | 0.04 [-0.07, 0.15] | 0.8 | 8 | 0.463 | ns | 0.49 [-1.00, 1.97] |
| Average (PRO pigs) | KM | $mL/cm^3$ | Vt | 5 | 5 | 0.753 | 8.3E+114 | -8e+114 [-3e+115, 1e+115] | -1.0 | 8 | 0.347 | ns | -0.63 [-2.14, 0.87] |

| | | | | | | | | | | | | | |
|---|---|---|---|---|---|---|---|---|---|---|---|---|---|
| Average (KD pigs) | KM | μL·cm$^{-3}$·h$^{-1}$ | K1 | 5 | 5 | 114.76 | 133.74 | -18.99 [-86.16, 48.18] | -0.7 | 8 | 0.533 | ns | -0.41 [-1.89, 1.07] |
| Average (KD pigs) | KM | h$^{-1}$ | k2 | 5 | 5 | 0.155 | 0.200 | -0.05 [-0.20, 0.11] | -0.7 | 8 | 0.508 | ns | -0.44 [-1.92, 1.04] |
| Average (KD pigs) | KM | mL/cm$^3$ | Vt | 5 | 5 | 4.7E+110 | 0.743 | 5e+110 [-6e+110, 2e+111] | 1.0 | 8 | 0.347 | ns | 0.63 [-0.87, 2.14] |
| Average (PRO+KD) | KM | μL·cm$^{-3}$·h$^{-1}$ | K1 | 5 | 5 | 100.47 | 120.14 | -19.68 [-81.06, 41.70] | -0.7 | 8 | 0.481 | ns | -0.47 [-1.95, 1.02] |
| Average (PRO+KD) | KM | h$^{-1}$ | k2 | 5 | 5 | 0.122 | 0.180 | -0.06 [-0.20, 0.08] | -1.0 | 8 | 0.366 | ns | -0.61 [-2.11, 0.89] |
| Average (PRO+KD) | KM | mL/cm$^3$ | Vt | 5 | 5 | 1.7E+109 | 0.739 | 2e+109 [-2e+109, 5e+109] | 1.0 | 8 | 0.347 | ns | 0.63 [-0.87, 2.14] |

### Table S4. End truncated KM comparisons (anesthesia groups pooled)

| Data | k2 parameter | Parameter | Unit | Pairwise comparison | N | Mean K1 | Mean K1 (6h KM) | Mean difference (K/D-PRO) [95% CI] | T-statistic | DF | p.value | Significance | Cohens' d [95% CI] |
|---|---|---|---|---|---|---|---|---|---|---|---|---|---|
| KM | free k2 | K1 | μL·cm$^{-3}$·h$^{-1}$ | 2h vs 6h | 10 | 89.20 | 108.82 | -19.62 [-29.96, -9.28] | -4.29 | 9 | 0.0020 | ** | -0.84 [-1.33, -0.36] |
| KM | free k2 | K1 | μL·cm$^{-3}$·h$^{-1}$ | 3h vs 6h | 10 | 95.46 | 108.82 | -13.36 [-19.82, -6.90] | -4.68 | 9 | 0.0012 | ** | -0.56 [-0.82, -0.29] |
| KM | free k2 | K1 | μL·cm$^{-3}$·h$^{-1}$ | 4h vs 6h | 10 | 98.76 | 108.82 | -10.06 [-15.89, -4.22] | -3.90 | 9 | 0.0036 | ** | -0.42 [-0.65, -0.18] |
| KM | free k2 | K1 | μL·cm$^{-3}$·h$^{-1}$ | 5h vs 6h | 10 | 103.79 | 108.82 | -5.03 [-8.17, -1.88] | -3.62 | 9 | 0.0056 | ** | -0.20 [-0.32, -0.08] |
| KM | k2 = 0 | K1 | μL·cm$^{-3}$·h$^{-1}$ | 2h vs 6h | 10 | 86.69 | 78.01 | 8.69 [-2.03, 19.40] | 1.83 | 9 | 0.0998 | . | 0.49 [-0.10, 1.08] |
| KM | k2 = 0 | K1 | μL·cm$^{-3}$·h$^{-1}$ | 3h vs 6h | 10 | 89.91 | 78.01 | 11.91 [4.86, 18.96] | 3.82 | 9 | 0.0041 | ** | 0.64 [0.25, 1.02] |
| KM | k2 = 0 | K1 | μL·cm$^{-3}$·h$^{-1}$ | 4h vs 6h | 10 | 87.34 | 78.01 | 9.33 [5.67, 13.00] | 5.76 | 9 | 0.0003 | *** | 0.54 [0.33, 0.75] |
| KM | k2 = 0 | K1 | μL·cm$^{-3}$·h$^{-1}$ | 5h vs 6h | 10 | 82.87 | 78.01 | 4.87 [3.29, 6.44] | 6.98 | 9 | 0.0001 | *** | 0.36 [0.25, 0.47] |
| KM | k2 = 0.13 | K1 | μL·cm$^{-3}$·h$^{-1}$ | 2h vs 6h | 10 | 89.51 | 103.80 | -14.29 [-24.74, -3.85] | -3.10 | 9 | 0.0128 | * | -0.79 [-1.40, -0.18] |
| KM | k2 = 0.13 | K1 | μL·cm$^{-3}$·h$^{-1}$ | 3h vs 6h | 10 | 98.11 | 103.80 | -5.69 [-11.80, 0.42] | -2.11 | 9 | 0.0646 | . | -0.33 [-0.67, 0.01] |
| KM | k2 = 0.13 | K1 | μL·cm$^{-3}$·h$^{-1}$ | 4h vs 6h | 10 | 102.70 | 103.80 | -1.10 [-4.24, 2.04] | -0.79 | 9 | 0.4478 | ns | -0.06 [-0.22, 0.10] |
| KM | k2 = 0.13 | K1 | μL·cm$^{-3}$·h$^{-1}$ | 5h vs 6h | 10 | 103.59 | 103.80 | -0.21 [-1.77, 1.35] | -0.30 | 9 | 0.7725 | ns | -0.01 [-0.11, 0.08] |
